# Decision Confidence Neuron in Echo State Network for Continual Evaluation of EEG Motor Imagery Classification Quality

**DOI:** 10.64898/2026.08.31.748424

**Authors:** Eva Lemoine, Brendan Lenfesty, Umesh Kumar Naik Mudavath, Saugat Bhattacharyya, KongFatt Wong-Lin

**Affiliations:** Faculty of Life Sciences, Bioinformatics Engineering Track, University of Poitiers, Poitiers Cedex 9, France; Intelligent Systems Research Centre, School of Computing, Engineering and Intelligent Systems, Ulster University, Magee Campus, Derry~Londonderry, Northern Ireland, UK; Department of Electronics and Electrical Engineering, Indian Institute of Technology Guwahati, Guwahati, Assam, India

## Abstract

Echo state networks (ESNs) are efficient, neuro-inspired computational frameworks well suited to time-series data. However, ESN decision confidence is typically quantified in limited ways. We propose an explicit decision-confidence readout neuron, trained from decision readout outputs, to continuously monitor confidence as decisions form. In a simulated decision task, confidence activity increased with stimulus strength, linking greater discriminability to higher confidence. We then evaluated the model on EEG-based motor imagery classification, showing that confidence activity increased with decision accuracy and discriminated correct from error decisions, particularly in higher-performing participants, reflecting human-like metacognition. Overall, this approach enables continual monitoring of decision confidence, supporting more trustworthy ESN decisions, particularly in biomedical applications.

## Introduction

In comparison with deep learning, which is popular in modern artificial intelligence (AI) applications, reservoir computing (RC) is inspired by the recurrent connectivity in biological brain networks [1], [2]. Given current emphasis on energy efficiency and edge analytics, RC is more favourable in these respects as it is substantially more lightweight and computationally efficient than, for example, deep networks or other recurrent neural networks [1], [2].

A specific type of RC is the echo state network (ESN) [3], [4]. Generally, ESN makes use of its fixed heterogeneous recurrent connectivity (“reservoir”) and fading trace or “echo” of past input signals which can be “tapped” into via their output combinations at readout decision neurons – this simplifies the learning through simple and fast regression only at the readout stage while avoiding complex and costly (e.g. backpropagation) learning within the reservoir [1], [3]. Further, due to the reservoir, ESN can compute nonlinear temporal patterns through linear separation at the readout layer.

There are wide applications of ESN. An application is in time-series predictions and forecasting (e.g. financial markets, weather forecasting, energy grids) [5]. Another example is in signal processing and telecommunications [6]. Other examples include pattern recognition and healthcare, e.g. speech recognition, biomedical telemetry, gesture classification [7], [8], [9]. Exploiting not only its efficiency but the multiple timescale in the reservoir being similar to the multiple timescale (and frequencies) in brain activity [10], ESN has been successfully applied to brain/mental health and neurotechnology, including in brain-computer interface (BCI), on neurodegeneration, motor imagery, and seizure [11], [12], [13], [14].

Despite these successes of ESN applications, the evaluation of ESN’s decision quality is limited. In particular, to compute the decision confidence of ESNs, it is conventional to use simple point-estimate outputs [4], prediction intervals [15], or Bayesian inference [16]. Thus, real-time computing of human-like decision confidence for trustworthy ESN decisions is currently lacking. This is particularly concerning if the applications are in critical situations, e.g. causing safety risks for patients.

In contrast, in cognitive neuroscience, specific neurons and brain regions encoding decision confidence have been observed [17], [18], [19]. When decisions are easier, these neural activities will tend to be more enhanced. Importantly, as decision-making in the biological brain tends to integrate evidence over time before committing to a decision among alternatives [20], the confidence-encoding neural activities also temporally evolve, providing real-time monitoring of confidence as the decision unfolds, which can be computed prior to decision commitment.

In this paper, inspired by neuroscience of decision-making and metacognition [20], [21], [22], we will develop an explicit decision confidence computing neuron at the readout layer of ESN. This will be trained based on the decision readout neurons of the same ESN; harder separation between decision neuronal activities will lead to lower decision confidence neuronal activity. Model training to optimise the decision confidence neuronal output will be conducted at the readout layer. Our proposed approach will be evaluated using a simulated standard two-choice task and a motor-imagery electroencephalography (EEG) based BCI data.

## Methods

### Echo State Network with Decision Confidence

The ESN model comprises 200 recurrent neural units, two input units and three output units. The activity, r(t), evolves as:

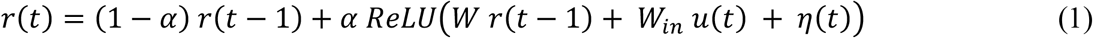

with *u*(*t*) denoting the input vector, *α*, the leak term and *η*(*t*) a rectified Gaussian noise. A ReLU neuronal activation function was applied to keep activity non-negative. *W*_*in*_, the input weights were sampled uniformly between 0 and 1 and normalised across channels to provide equal drive to the reservoir. *W*, the recurrent weights, were sampled from a zero-mean Gaussian distribution, with recurrent weights zeroed in accordance with the sparsity parameter, and the remaining weights were rescaled to some specified spectral radius.

A softplus nonlinearity was applied to the product of the output weights and activity to ensure non-negative values, interpretable as firing rate activity. The weights of the two decision readout neurons were trained with ridge regression and represent one decision or the other. A decision is made when whichever of the decision neuronal activity first crossed a prescribed threshold of 5 = 1.2. Decision time is hence computed from input stimulus onset to decision threshold crossing.

An additional readout neuron was trained to assess the decision confidence with its own set of weights. Again, it was trained separately by non-negative ridge regression to predict the absolute difference in decision evidence between the two decision readout neurons. Confidence readout therefore represents the separation of the two competing decision neuronal activities, which tend to separate over time akin to neural evidence accumulation between options during decision formation in decision neuroscience [20].

### Model Training and Testing in Simulated 2-Choice Task

The model was first tested on two noisy inputs for a standard two-choice discrimination task, often used in perceptual decision-making literature [20], [23]. The model parameters were initially tuned to this task, where the network must decide whether the stimulus is stronger for one input or the other, based on the accumulated evidence over time. The third readout neuron was trained to compute decision confidence continuously while the evidence was accumulated towards a threshold. This simulated task was set up to validate the network architecture and function wherein the ground truth of the decision (based on the presented input stimulus difference) was known, before applying to EEG data, where it can be more challenging to interpret decision confidence.

The simulated task trial had an initial 500 *ms* of minimal baseline activity with background noise of amplitude 0.05 before the onset of two noise-corrupted step-function stimulus inputs that lasted for 900 *ms*, and can be described, without loss of generality that decision 1 is the correct decision, by:

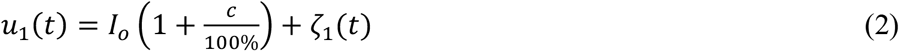

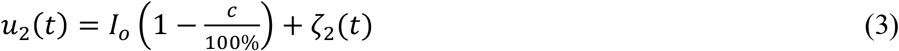

with independent Gaussian (rectified) noise *ζ*_1_ and *ζ*_2_ of amplitude 0.625 on top of baseline input strength *I*_0_ = 0.5, and normalised input strength *c* B {0, 4, 8, 16, 32, 64} %. Note that the deterministic input difference is 2*c*.

For this task, we set the model parameter values such that *α* = 0.3, noise amplitude was 0.295, network sparsity was 80%, and spectral radius was 0.8. A spectral radius below 1 was required throughout to satisfy the echo-state property, ensuring stable, non-divergent reservoir dynamics [3]. A reservoir size of *I* = 200 units were chosen manually as it was big enough to enable emergent population-level dynamics but remained computationally lightweight for further application, as used in real-time BCI.

The two decision readout neurons were trained by ridge regression on 1000 trials per input difference per direction of the reservoir activity. The confidence readout neuron was trained separately using a further 100 trials per input strength; the smaller the strength (i.e. signal), the harder was the task.

The model was tested with decision accuracy and decision time. Their performances were evaluated based on decision neuroscience literature [20], namely, via decision accuracy vs. (absolute) stimulus input strength (psychometric graph), and decision time vs. (absolute) input strength (chronometric graph). This testing took place across 3000 trials. Decision confidence was evaluated for its variation with stimulus input strength to test whether the readout scales with overall stimulus discriminability regardless of the decision made. Separation between correct and error trials at matched input strength was also assessed via statistical analysis and area under the ROC curve (AUC).

### BCI Data Description and Preprocessing

For validation on actual data, the ESN with decision neuron was validated on an open EEG based BCI data for potential real-time deployment. In particular, we used the BCI Competition IV Dataset 2b [24], a public dataset used for benchmarking motor imagery classification for right/left hand EEG and openly available via the original study [24] (https://www.bbci.de/competition/iv/). The data contains 9 participants, 3 sessions, and 2 motor-imagery classes: left hand (LH) and right hand (RH). We used the 3 labelled sessions (01T, 02T, and 03T) for each participant, and we retained the 3 EEG channels (C3, Cz, and C4) (Fig. 1A) to bridge towards practical applications in which the number of EEG electrodes are more limited.

**Figure 1.**
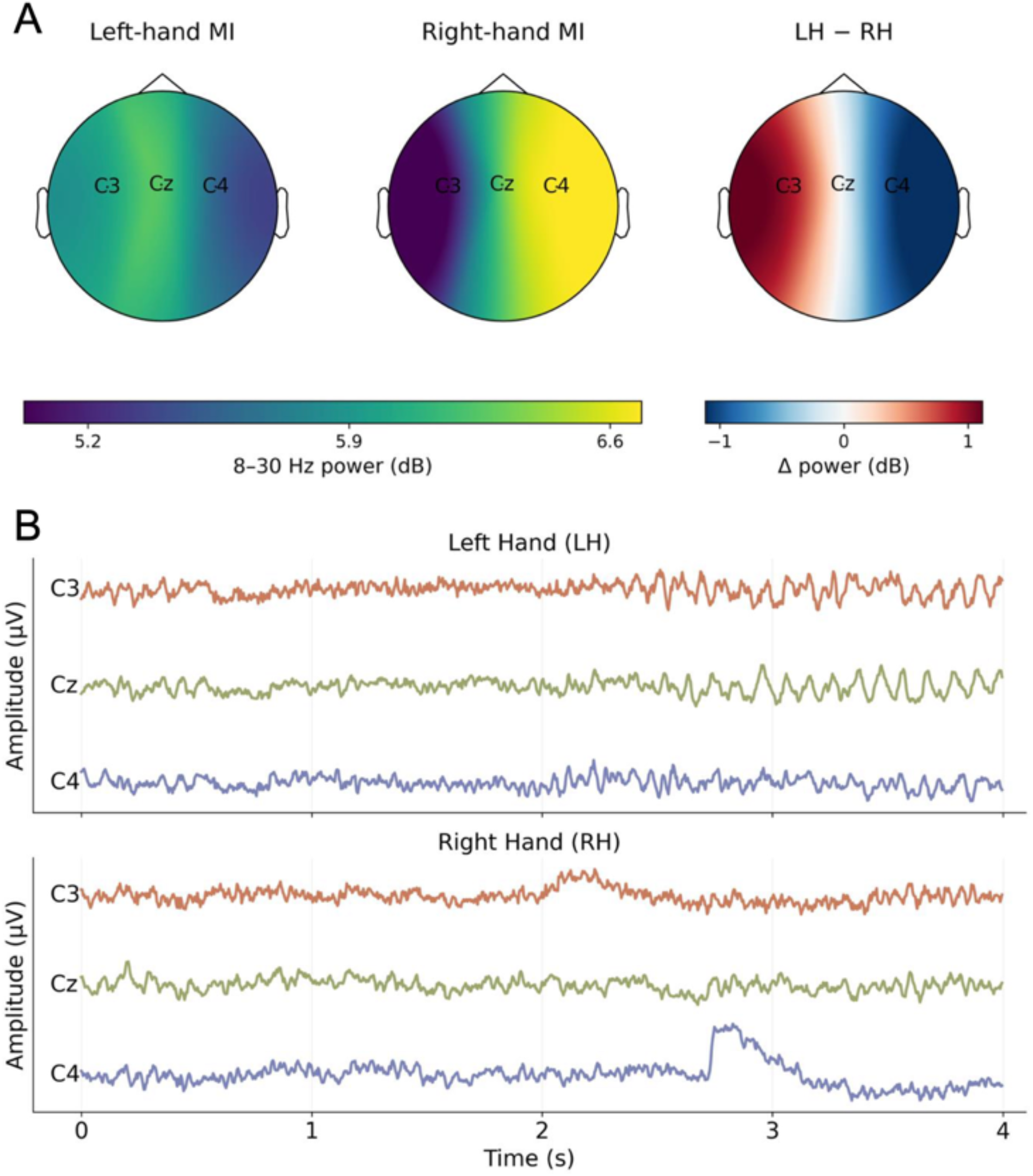
Preprocessed EEG motor imagery data. (A) Grand-average topographical maps of EEG. Left-hand (left) and right-hand (middle) motor imagery, and their class-wise difference (right). (B) Sample left-hand (LH) (top) and right-hand (RH) (bottom) motor imagery across channels C3, Cz, and C4 from Subject 1, session 1.

The original recordings were sampled at 250 *Hz*, band-pass filtered between 0.5 and 100 *Hz*, and recorded with a 50 *Hz* notch filter enabled. As part of our preprocessing, trials marked as artefacts in the original dataset were excluded, and each trial was segmented into a 4-second cue-locked epoch, giving 1,000 samples per channel. The final dataset contained 3,026 epochs, comprising 1,514 LH and 1,512 RH trials, resulting in an approximately balanced 2-class dataset and providing a larger benchmark for evaluating the model compared with the initial 4-class dataset used in our initial testing. Sample preprocessed of the 3-channel EEG data is depicted in Fig. 1B.

### Training and Testing on BCI Data

When the ESN model was trained to classify the BCI Competition IV dataset 2b, sessions 1 and 2 were used for training and validation, and session 3 was untouched and only used for testing the trained model. The electrodes used were mu and beta band filtered (8 − 13 *Hz*, 13 − 30*Hz*), the amplitude envelopes converted using Hilbert transform, with normalisation to sum to 1 across channel for each timestep, similar to the input in the simulated task. We expanded the reservoir to take 4 inputs and 6 inputs for comparison with mu+beta band pass classification. The 4 inputs were C3/C4 channels for both mu and beta bands, and with 6 inputs expanding to C3/C4/CZ for the same bands. The 4-input model outperformed the 6-input model, hence the former was used for subsequent analysis. This could be because C3 and C4 event related desynchronisation for left vs right hand imagery was more directly lateralised between the two hemispheres than the midline channel, Cz.

In this task, the hyperparameters (spectral radius was 0.8, and leak rate was 0.1, noise was 0, sparsity was 0.5, and *N* = 400) were initially tuned using a grid search for sessions 1 and 2 only; ridge strength was also chosen based on sessions 1 and 2 only. The tuned model was then trained on pooled sessions 1 and 2, and then evaluated on session 3. Chance level was not assumed and was worked out empirically with permutation tests. Band-power features were used as reference for model performance as per its standard use for such a task (Fig. 1A).

The confidence readout neuron was fitted by non-negative ridge regression on training sessions 1 and 2 only, using the tuned input reservoir. As there is no ground truth for confidence signal for EEG data, unlike the simulated task, the readout was evaluated against trial correctness on session 3. Confidence was non-normally distributed for most of the 9 participants (Shapiro-Wilk test, 7 of 9 non-normally distributed). Consequently, discrimination between correct and error trials was assessed with appropriate statistical tests (ROC curve (AUC) Mann-Whitney U test). Replication was then tested across the group with Wilcoxon signed rank test for all 9 participants; AUC values against chance, with a binomial sign test on the direction of the effect. Secondary analysis used participant z-scored point biserial correlation for comparability to prior literature. Trial correctness was summarised as accuracy within confidence quartiles, with confidence z-scored per participant and then pooled to allow comparison for less to more confidence bins without between participant accuracy differences muddling results. The analytical workflow is summarised schematically in Fig. 2.

**Figure 2.**
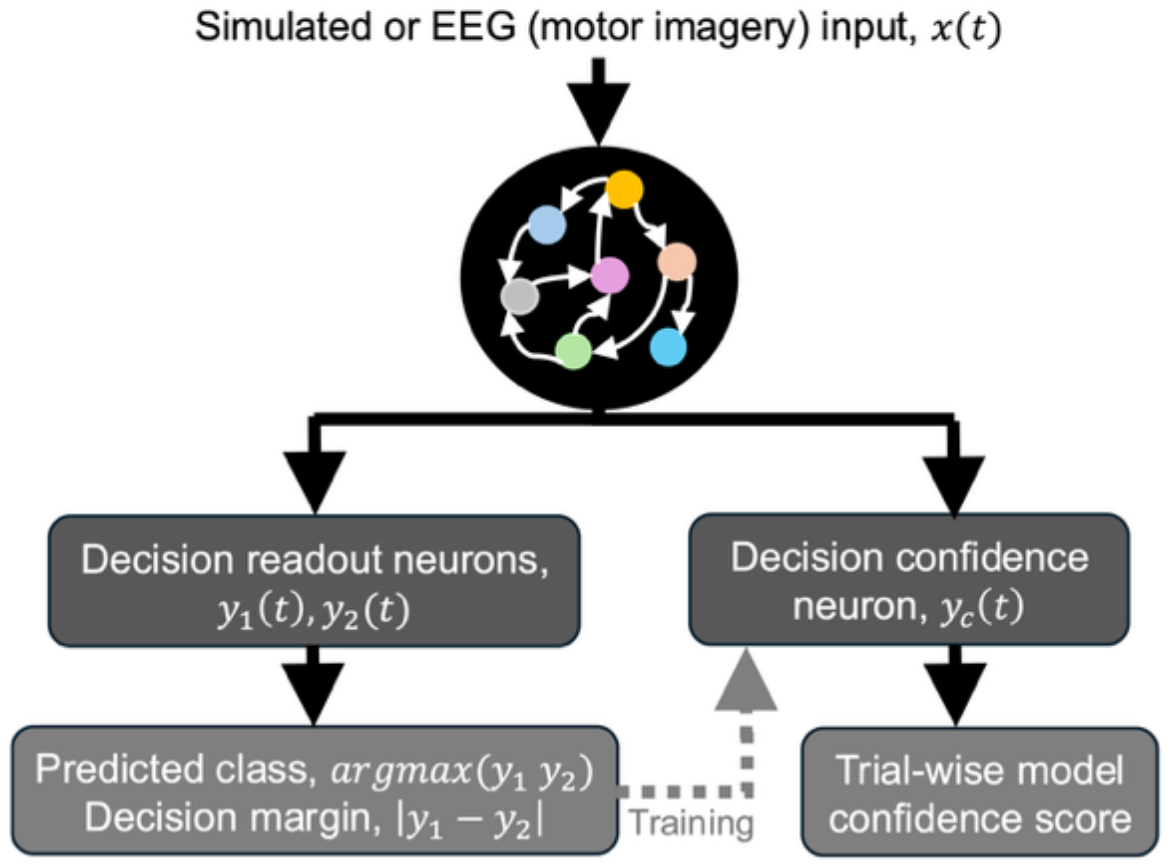
Workflow for training and evaluating the echo state network (ESN) with continual decision confidence readout. Fixed ESN’s reservoir (coloured joined nodes) provides shared recurrent state to 2 decision readouts and a confidence readout. Separation between decision outputs used as training target for confidence readout. During testing or validation, the model provides the predicted class together with trial-wise confidence estimate.

### Software and Hardware

Model implementation, training, and analysis were performed in Python 3.9 using Jupyter Notebook (Intel Core i7-14700HX, 32 GB RAM). The EEG signal preprocessing was performed using Python 3.8 with Google Colab. Source code will be made available upon publication.

## Results

### ESN Decision Neuron can Compute Discriminability in Simulated Stochastic Decision Task

The ESN’s decision neurons readily exhibited appropriate decision behaviours on the simulated two-choice task described in Section IIB. The psychometric graph (Fig. 3A) shows a rise towards 100% choice accuracy from chance level as input strength increases from ambiguous signal of 0%. The chronometric graph (Fig. 3B) illustrates a typical trend of decreasing decision time for correct trials from harder to easier decisions of increasing stimulus input strength. For error trials, if any, the decision times were generally slower than that of correct trials, similar to human-like decision-making trends [25].

**Figure 3.**
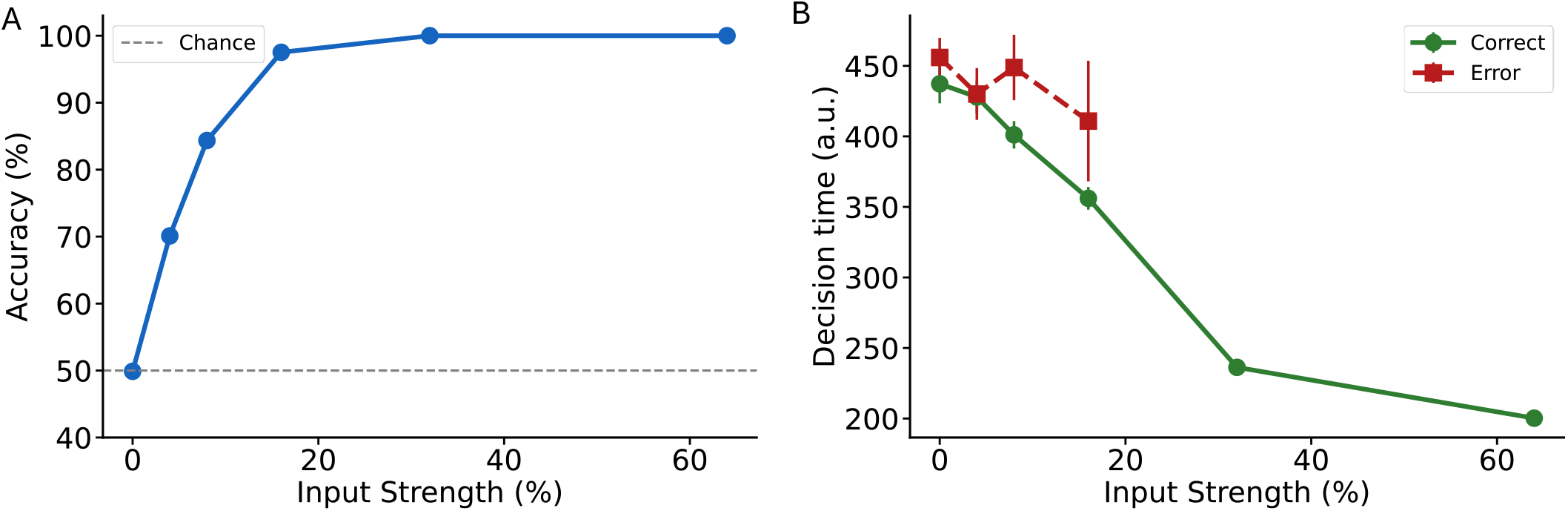
ESN decision behaviours in simulated task. (A) Psychometric graph: accuracy rises from chance (50%) at ambiguous stimulus (zero input difference) towards perfect performance with increasing input strength. (B) Chronometric graph: decision time decreases with input strength and is generally slower for error than correct trials.

The decision confidence neuronal activity, trained with ridge regression from the reservoir, generally predicted the difference between decision neuronal outputs at the decision threshold, tracking the strength of the evidence (Fig. 4A). Note that the output activity was minmax 9ormalized in Fig. 4 for visual purposes.

**Figure 4.**
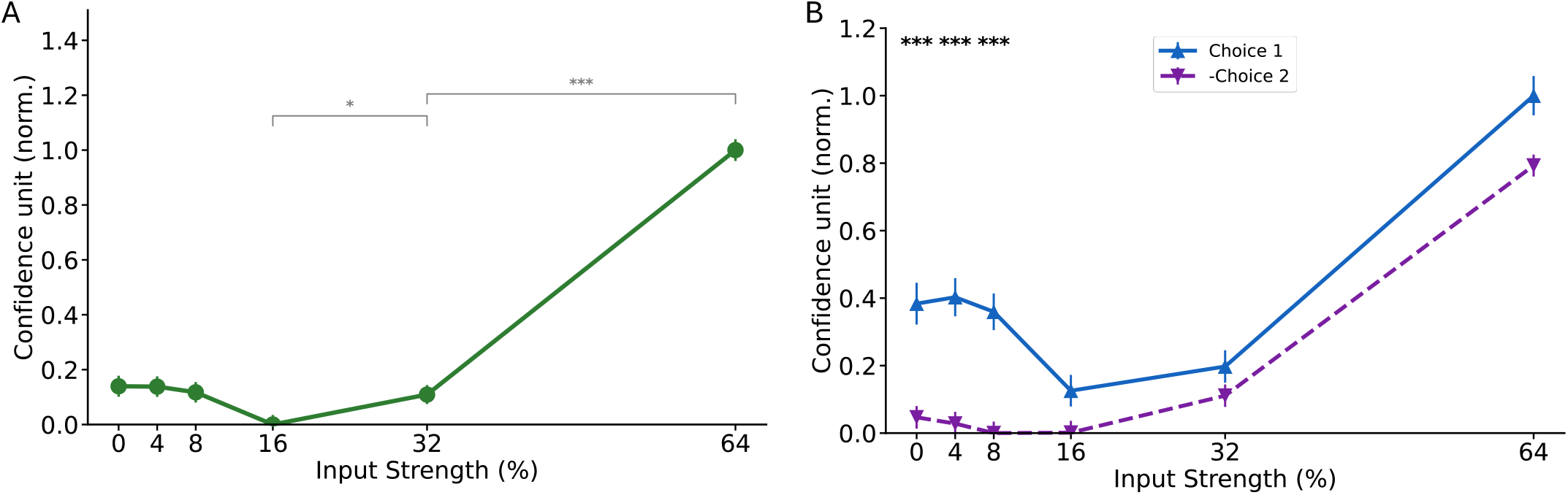
ESN confidence neuronal output in simulated task generally increases with easier task and agnostic to specific choices. (A) Confidence at decision, pooled across choice and outcome, increases monotonically with input difference (mean ± SEM, normalized; Jonckheere-Terpstra trend test, inset). (B) The same measure split by choice direction (mean ± SEM, normalized) shows a small but significant sign-dependent bias at low input difference (0–8%; asterisks, FDR-corrected), which is not significant at higher input difference (16–64%).

Statistical analysis was done on a trial-by-trial basis. Specifically, we could observe the decision confidence neuronal activity exhibiting an increase with input strengths after a certain intermediate value of ~16%. A Jonckheere-Terpstra Trend test indeed showed a significance of the increasing neuronal response to increasing input strength; this was further verified with a Mann-Whitney U test and corrected with Benjamini-Hochberg FDR for a significant difference between strengths of 16% and 32%, as well as between 32% and 64%.

Further, the decision confidence remained relatively agnostic to the input difference sign, i.e. the specific choice made (Fig. 4B). Statistical analysis for specific choices showed the network was biased with lower input strengths at ~16%. However, at high input strengths, the confidence neuron was not significantly biased towards either choice. This was verified with Mann-Whitney U between pairs for matched input strength and corrected with Benjamini-Hochberg FDR. Thus, the trends were similar to the observed neuronal encoding of decision confidence [17], [18], [21].

Despite the promising results so far shown by the decision neuron, the separation between correct and error trials was generally minimal (Fig. 5A), which was further validated using AUC (Fig. 5B). Thus, the decision confidence neuronal activity in this task seemed to reflect stimulus discriminability rather than the correctness of choice, though EEG data will enable further examination.

**Figure 5.**
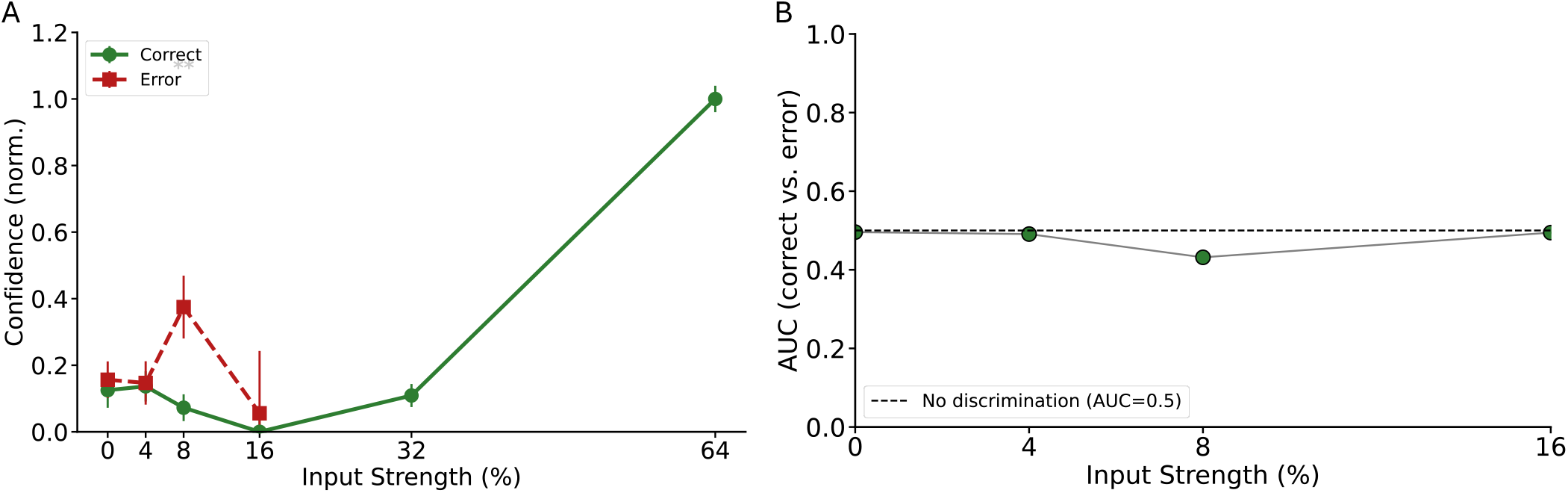
Decision confidence neuron in simulated task does not reliably discriminate correct from error choices. (A) Confidence at decision, for correct and error trials, across input-difference levels (mean ± SEM; normalized). (B) Area under the ROC curve (AUC) quantifying correct-vs-error separation, restricted to input-difference levels with a sufficient number of error trials (≤ 16%; grey markers indicate levels excluded for insufficient error trials).

### ESN Decision Neuron Can Compute Metacognition in EEG-BCI Motor Imagery Classification

The initial tests (using session 3) of the ESN model involved comparing the 4-input model vs mu+beta-band classification (the standard approach). The mu+beta-band classification was 71% compared to the model with 72.3%, which thus had the best classification performance.

The model was further tuned from the original model parameters. A grid search was performed on the model and trained on session 1 while evaluated on session 2. The original model had an accuracy of 65.2% prior to tuning but subsequently improved to 67.5% from the tuned parameters (mentioned in Section II). The tuning evaluation on the model improved its accuracy to 72.6% compared to mu-band+beta-band classification of 71% (Fig. 6). We found that the classification accuracy varied consistently with participants reflecting signal quality rather than a limitation to the computational approaches (Fig. 6).

**Figure 6.**
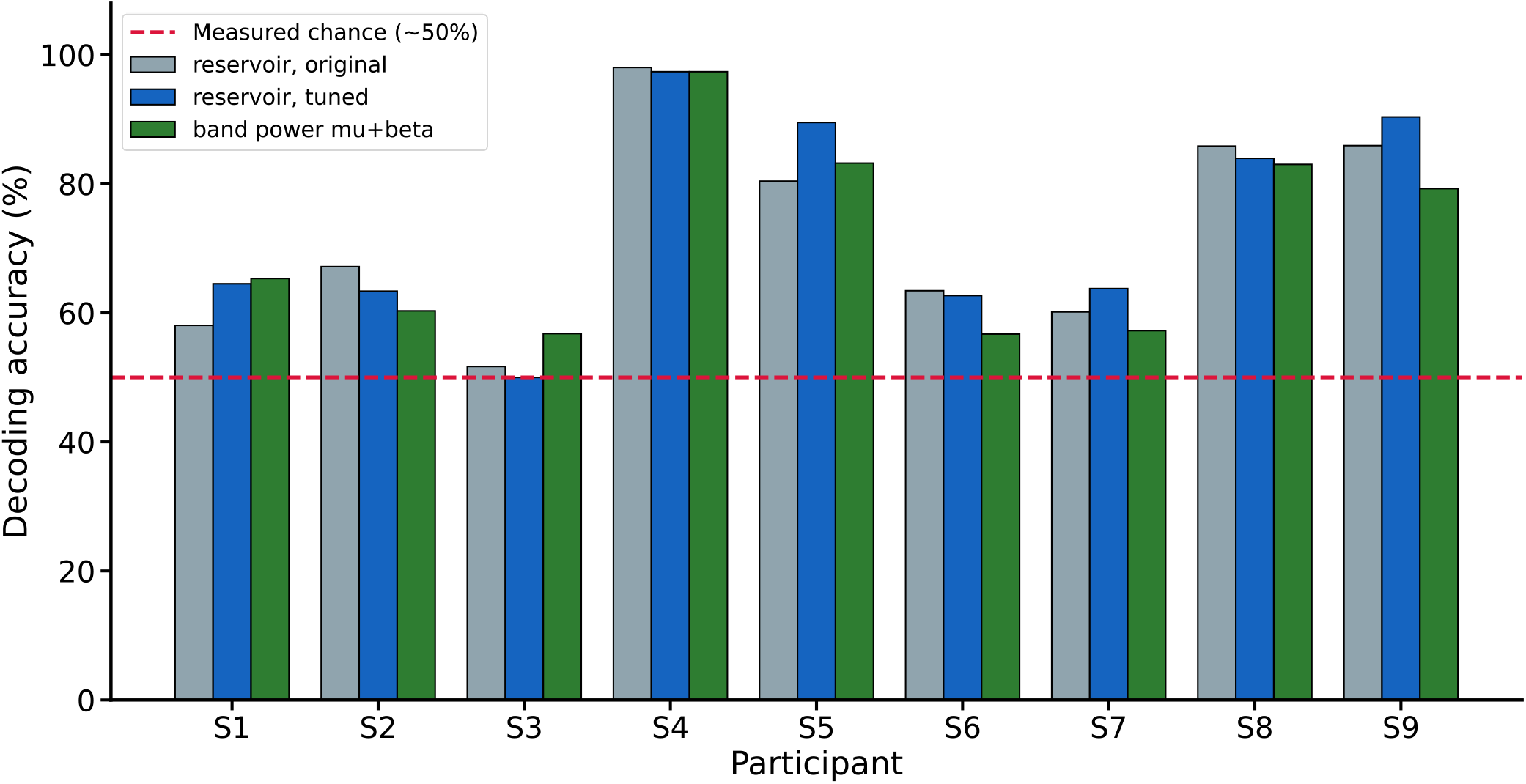
Decoding accuracy in EEG-BCI motor imagery task. (A) Accuracy per participant for the original (grey) and tuned (blue) 2-channel input ESNs, compared against the band-power reference feature (green) ESN. Red dashed line: measured chance level of 50%.

To test the confidence neuron’s ability to identify strong signal and correctness of decisions, we performed statistical analysis per participant with AUC and Mann-Whitney U test. We found the confidence neuron to classify correct from error choices above chance level in all 9 participants (mean AUC = 0.639, SD = 0.116), and individually significant from chance in 4 of 9 participants (Fig. 7, green). This was then tested across the group, and again, proved statistically significant against chance classification using the Wilcoxon signed rank test (*p* = 0.0020), and additionally with the standard binomial sign test (*p* = 0.0020). Due to participant S4 having only 3 error trials, we removed S4’s data and performed the tests again, resulting in the mean AUC= 0.607, still being greater than chance while statistical significance remained for both group-wise comparisons (Wilcoxon *p* = 0.0039, binomial sign *p* = 0.0039). A standard z-scored point biserial correlation gave a similar picture (*r* = +0.113, permutation *p* < 0.001).

**Figure 7.**
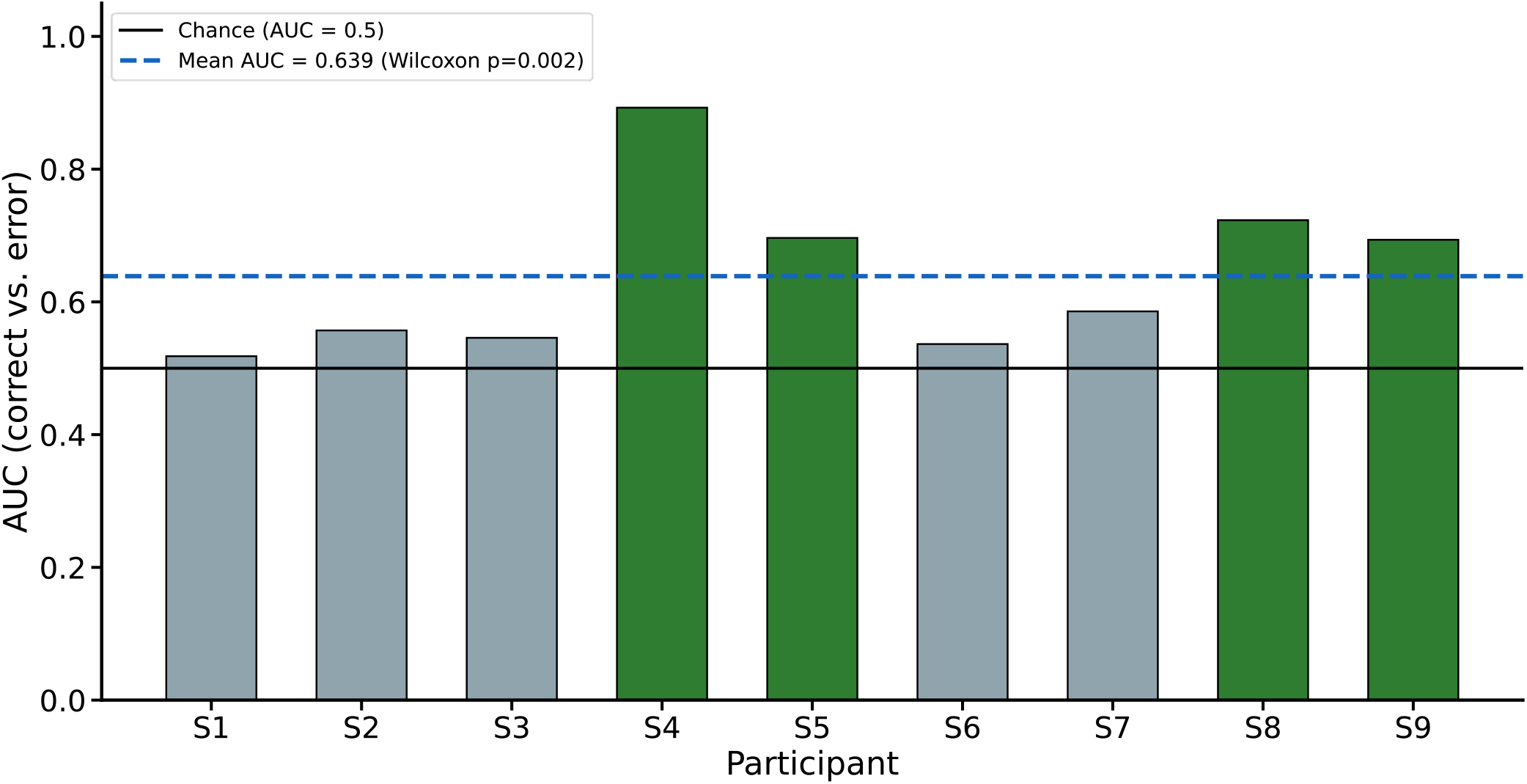
Decision confidence discriminates correct from error choices in EEG-BCI motor imagery data. Area under the ROC curve (AUC), per participant, quantifying separation between confidence on correct versus error trials (green bars: individually significant participants, Mann-Whitney U, *p* < 0.05). Dashed black line: chance (AUC= 0.5); dashed blue line: mean AUC across participants (Wilcoxon signed-rank test against chance).

Having established that the confidence neuron could encode choice correctness, a measure of metacognitive processing, could it also encode higher confidence for the more accurate trials? Confidence neuronal outputs were pooled for a given trial per participant, and we z-scored them to enable comparability across participants. The participants’ trials were then split into the four quartiles (Q’s) and combined to see if the quartiles representing higher confidence also had higher accuracy. Indeed, the results showed a monotonic increase from low to high confidence output as decision accuracy increased (Fig. 8). Overall, this demonstrates that the decision confidence neuron could readily track not only choice correctness but also choice accuracy, suggesting a potential use case for ESN.

**Figure 8.**
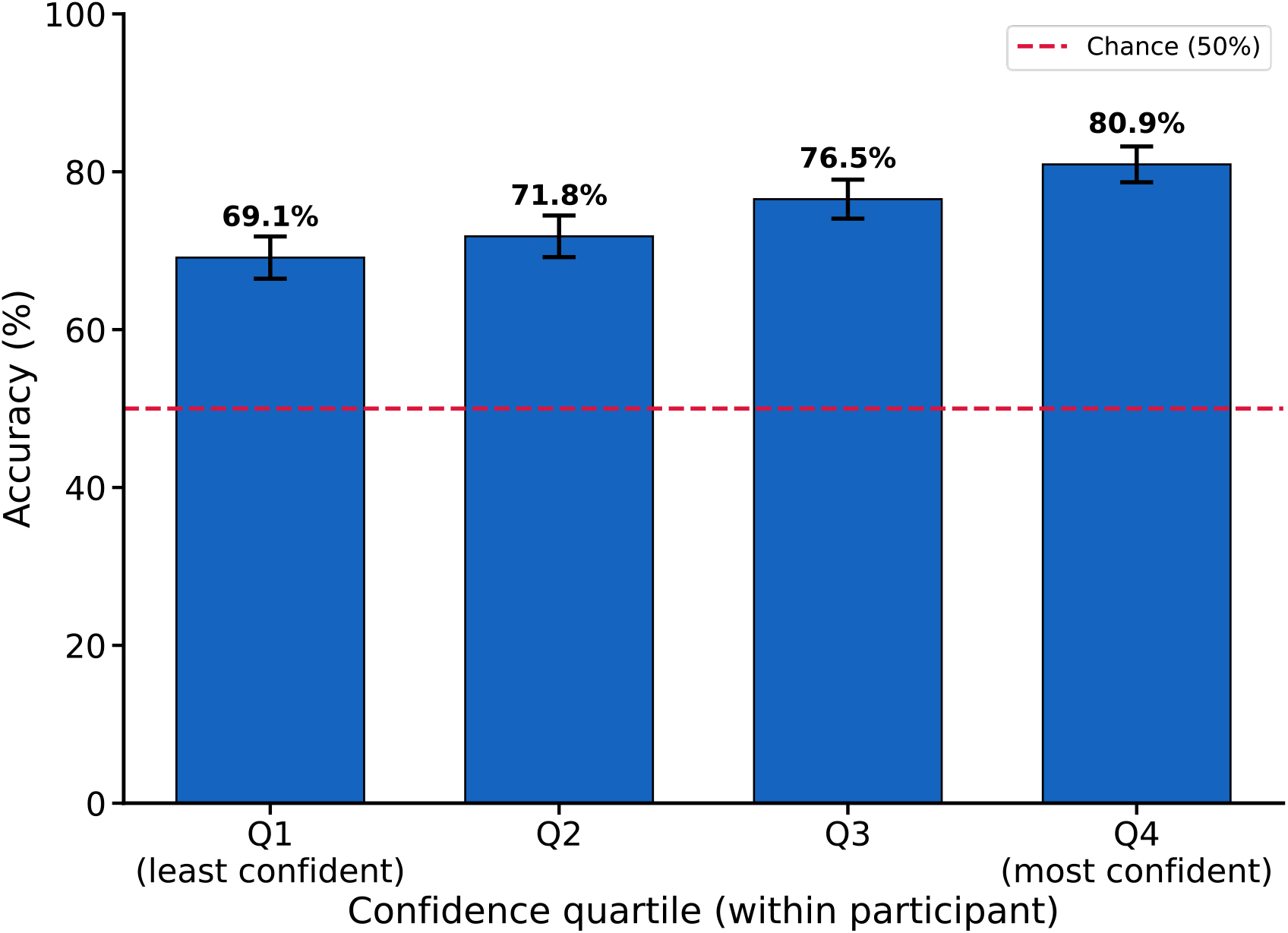
Decision accuracy scales with confidence quartile in EEG-BCI motor imagery task. Decoding accuracy within each confidence quartile, pooled across participants after within-participant z-scoring (mean ± SEM). Accuracy generally rises from the least (Q1) to the most (Q4) confident quartile.

## Conclusion

In this work, we have built on existing ESN modelling by incorporating an explicit decision readout neuron to continuously monitor decision confidence (Fig. 1). We first tested the modified ESN on a simulated stochastic two-choice task. We found that although the ESN’s decision neuron could compute signal discriminability and overall decision accuracy, it had difficulty computing decision correctness. This could be due to the overall stationary of the signal in the simulated task, leading to decision neuronal dynamics similar to a basic drift-diffusion model which could not distinguish confidence between correct from error choices due to their equal mean decision times [26]. Further analysis would be required to confirm this speculation.

The ESN’s decision neuron was later applied to an actual EEG-BCI based motor imagery task. In this use case, the ESN’s decision neuron was able to readily distinguish not only decision accuracy, but also decision correctness, which is together similar to human-like metacognition [22].

Taken together, we have successfully incorporated a decision neuron in the ESN to monitor the model’s decision confidence in real-time, while improving classification from the standard mu+beta band classification approach. This adds on top of the already efficient existing ESN model. In terms of applications, we believe that the decisions made by ESN can now be trusted more (e.g. when confidence is high). Moreover, as the ESN confidence exhibits human-like metacognition, human end users can better understand the decisions made. Finally, when decision confidence is low, it allows human end users to set appropriate safety threshold to terminate impending potentially dangerous decisions [27], [28], or disallow further machine execution e.g. in BCI-controlled wheelchair [29].

## Acknowledgement

This work was supported by Health and Social Care Research and Development (STL/5540/19) and Medical Research Council (MC_PC_20020), and The Royal Society (IEC\NSFC\252805). E.L. gratefully acknowledges the financial support from the European Union’s Erasmus+ Programme.

## Notes

**Conflict of interest statement:** The authors declare no competing interests.

### Competing Interest Statement

The authors have declared no competing interest.

